# From Genes to Pathways: A Curated Gene Approach to Accurate Pathway Reconstruction in Teleost Fish Transcriptomics

**DOI:** 10.1101/2024.09.23.614382

**Authors:** Marcela Herrera, Stefano Vianello, Laurie Mitchell, Zoé Chamot, Catherine Lorin-Nebel, Natacha Roux, Laurence Besseau, Yann Gibert, Vincent Laudet

## Abstract

Interpreting the vast amounts of data generated by high-throughput sequencing technologies can often present a significant challenge, particularly for non-model organism. While automated approaches like GO (Gene Ontology) and KEGG (Kyoto Encyclopedia of Genes and Genomes) enrichment analyses are widely used, they often lack specificity for non-model organisms. To bridge this gap, we present a manually curated gene list tailored for teleost fish transcriptomics. This resource focuses on key biological processes crucial for understanding teleost fish physiology, development, and adaptation, including hormone signaling, various metabolic pathways, appetite regulation, digestion, gastrointestinal function, vision, ossification, osmoregulation, and pigmentation. Developed through collaborative efforts of specialists in diverse fields, the list prioritizes genes with established roles in teleost physiology, experimental evidence, and conservation across species. This curated list aims to provide researchers with a reliable starting point for transcriptomic analyses, offering a carefully evaluated set of genes relevant to current research priorities. By streamlining the process of gene selection and interpretation, this resource supports the broader teleost fish research community in designing and analyzing studies that investigate molecular responses to developmental and environmental changes. We encourage the scientific community to collaboratively expand and refine this list, ensuring its continued relevance and utility for teleost fish research.

## Context

The advent of high-throughput sequencing and gene profiling technologies has revolutionized biological research, generating vast amounts of data that present both immense potential and significant challenges for meaningful interpretation (Khatri et al., 2012). Functional-level analysis has emerged as a powerful approach for managing this complexity by consolidating thousands of genes into smaller, biologically relevant sets and identifying active pathways that vary between conditions (Khatri et al., 2012). Two key resources in this landscape, the Gene Ontology (GO) project (Ashburner et al., 2000) and the Kyoto Encyclopedia of Genes and Genomes (KEGG) database (Ogata et al., 1998), have become invaluable references for interpreting high-throughput data within a broader biological context. These platforms provide comprehensive frameworks to summarize gene functions across multiple levels, from molecular interactions to ecosystem-level processes. GO term enrichment analysis offers a high-level overview of biological processes, cellular locations, and molecular functions associated with gene sets. On the other hand, KEGG pathway analysis provides more detailed insights into specific molecular mechanisms and pathways, supported by visual representations that make it easier to comprehend complex biological systems and their interactions. Thus, integrating both approaches improves our ability to interpret large-scale gene expression data, identify key biological processes, and ultimately advance our understanding of the molecular basis of life. These complementary approaches leverage extensive curated information to provide broad coverage of biological processes, molecular functions, and pathways across diverse organisms. Their power lies in their efficient data processing, standardized terminologies, and robust statistical methods, ensuring consistent interpretations and cross-study comparability (Blake & Harris, 2003; Kanehisa et al., 2012).

These automated approaches, while efficient, are not without drawbacks. reference databases are constantly evolving, with frequent updates and changes. For example, the Gene Ontology (https://geneontology.org/) undergoes regular revisions where new GO terms are created, some are merged, and others are made obsolete (Fig. 1). The number of GO terms by category (i.e., biological process, cellular component, or molecular function) also changes over time, as does the number of annotations by evidence type, which varies among different organisms. These dynamics can introduce the unfortunate drawback of complicating the immediate comparability between enrichment results across species and over time. This can pose challenges when comparing new analyses with previous literature or when attempting to draw cross-species comparisons. In contrast, approaches that maintain consistent terms over time, with only new genes being added, can facilitate easier comparison and interpretation of results across studies and species. Moreover, GO terms and KEGG pathways may lack specificity for precise biological questions, and database annotations can introduce bias, especially for non-classical model species or pathways. In our experience, GO- and KEGG-based enrichment analyses often fall short in fully capturing the underlying biology of the experimental intervention (whether it involves gene overexpression, gene mutation, or chemical exposure), the experimental context, or tissue-specific variations. Complex processes can be oversimplified in pre-defined categories that fail to encompass the subtleties of specific biological processes or evolutionary patterns. Recently discovered or hypothesized interactions may be absent from databases, and interpreting up- and down-regulation within pathways can still be challenging.

**Figure 1.**
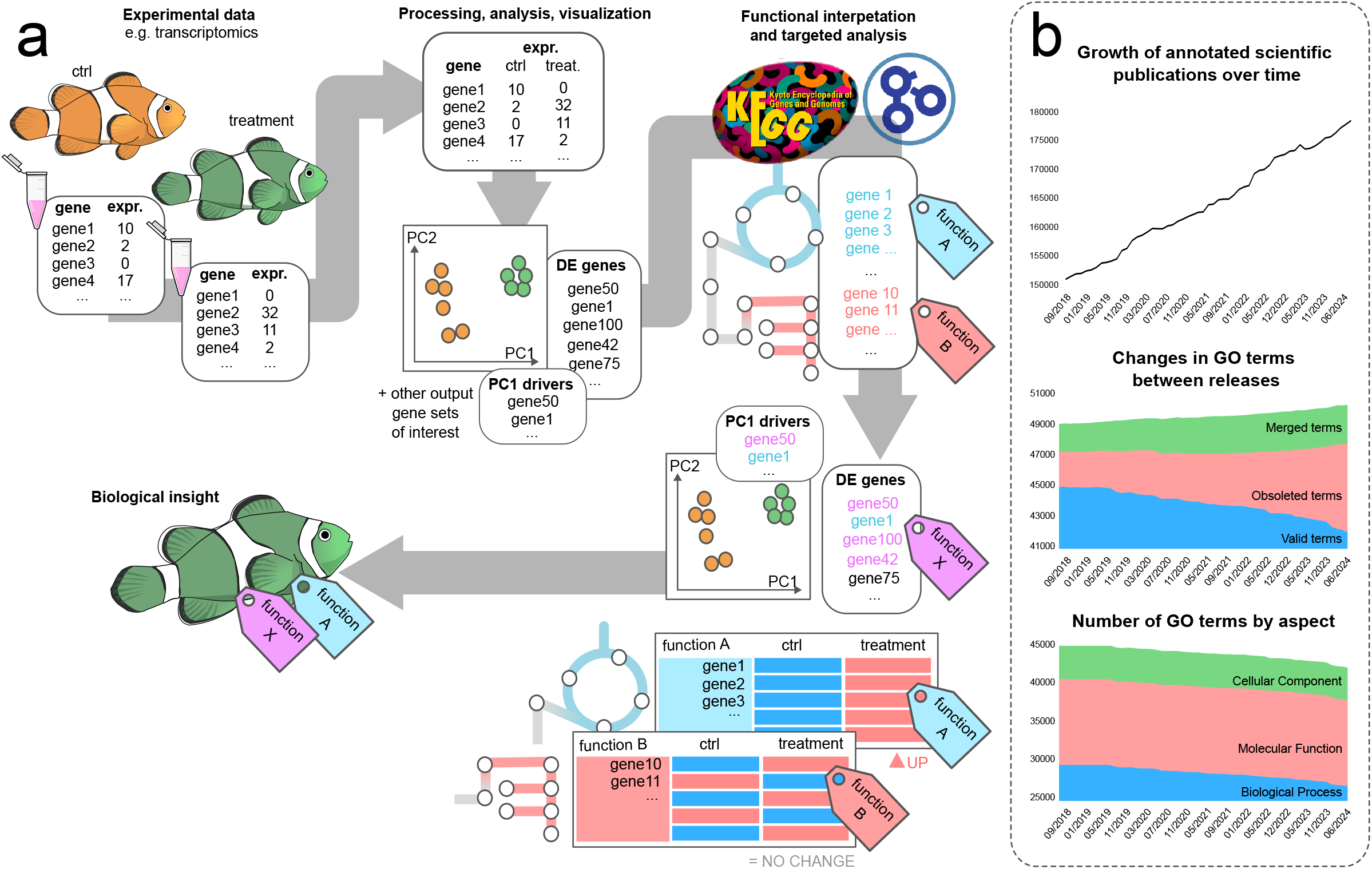
a) The generation, processing, analysis, and visualization of transcriptomic data. Functional analysis using Gene Ontology (GO) and KEGG pathways provide meaningful biological insights. **b)** The increasing number of scientific publications utilizing GO term annotations over time. The Gene Ontology undergoes regular updates, with the creation of new terms, merging of existing ones, and obsolescence of others. Consequently, the number of GO terms by category (biological processes, molecular function, and cellular component) varies over time. Data were retrieved from the Gene Ontology website and plotted by month and year of publication, covering the period from September 2018 to June 2024.

FishEnrichr, developed in 2018 as part of the ModEnrichr toolkit (Kuleshov et al., 2019), addresses some limitations of general databases in teleost research by providing a tool specifically tailored for zebrafish (*Danio rerio*). While this tool offers valuable automated annotations for interpreting fish transcriptomic data, we believe that manually curated gene sets provide several key advantages. Unlike FishEnrichr’s gene set libraries, which are based on data from 2018 and 2019, manually reconstructing gene pathways offers unique advantages despite requiring extensive literature review and being more limited in scope compared to established databases. This approach allows for greater specificity to the research question at hand and facilitates the integration of multiple data types and expert knowledge, leading to a more comprehensive and nuanced understanding of the biological system under study. Furthermore, manual reconstruction can also help identify knowledge gaps, potentially guiding future research directions. The study by Lorin et al. (2018) on teleost fish-specific retention of pigmentation genes provides a compelling example of the benefits of manual gene set curation. The authors created a highly accurate dataset by verifying GO annotations, integrating information from multiple sources, and applying a strict, context-specific definition of pigmentation genes (Lorin et al., 2018). They focused on neural-crest derived pigment cells differentiation across vertebrate species, excluding other types of pigment cells such as those of the retinal pigmented epithelium. To establish their pigmentation genes dataset, Lorin et al. initially retrieved genes annotated under the GO term “pigmentation” in vertebrates. However, they capitalised on manual curation to validate these automatic annotations against their specific definition of pigmentation genes. This process ensured that only genes directly involved in pigment cell differentiation were included. For instance, a gene like *superoxide dismutase 2* (*sod2*) was excluded despite its initial GO annotation as it did not pass their strict criteria (in this case *sod2* is also known for causing red blood cell damage). The authors also incorporated additional pigmentation genes from previous literature, categorizing them based on specific functions. This comprehensive approach not only curated a newer and more robust dataset of pigmentation genes but also highlighted the need of manual curation to refine functional categories beyond automated methods like GO annotation alone. It ensured the accuracy and relevance of their gene dataset, which is crucial for advancing our understanding of evolutionary and functional aspects of pigmentation in vertebrates.

Manual reconstruction is not without its challenges. It is typically time-consuming and labor-intensive, which can be a significant burden in large-scale studies. Recognizing this, we have been building upon the initial efforts of Roux et al. (2023), who characterized the transcriptomic landscape of metamorphosis in the false clownfish *Amphiprion ocellaris*. Thus, with the aim of streamlining teleost Eco-Evo-Devo research efforts and supporting the broader global scientific community, we are now sharing a carefully curated gene list focused on key biological processes crucial for understanding teleost fish physiology, development, and adaptation (Fig. 2, Fig. 3). We emphasize hormone signaling pathways, particularly thyroid hormones and corticoids, due to their pivotal roles in metamorphosis, growth, stress responses, and osmoregulation (Zwahlen et al., 2024). Metabolic pathways such as glycolysis, lactic fermentation, the citric acid (TCA) cycle, fatty acid β-oxidation, cholesterol biosynthesis, and lipid metabolism, are included to elucidate the complex biochemical processes that underpin cellular energy production and metabolic regulation (Fisher, 2001). Complementarily, appetite regulation genes are included to illustrate their role as key mediators between food intake and energy expenditure, influencing both short-term feeding behavior and long-term metabolic homeostasis (Conde-Sieira et al., 2018; Rønnestad et al., 2017).

**Figure 2.**
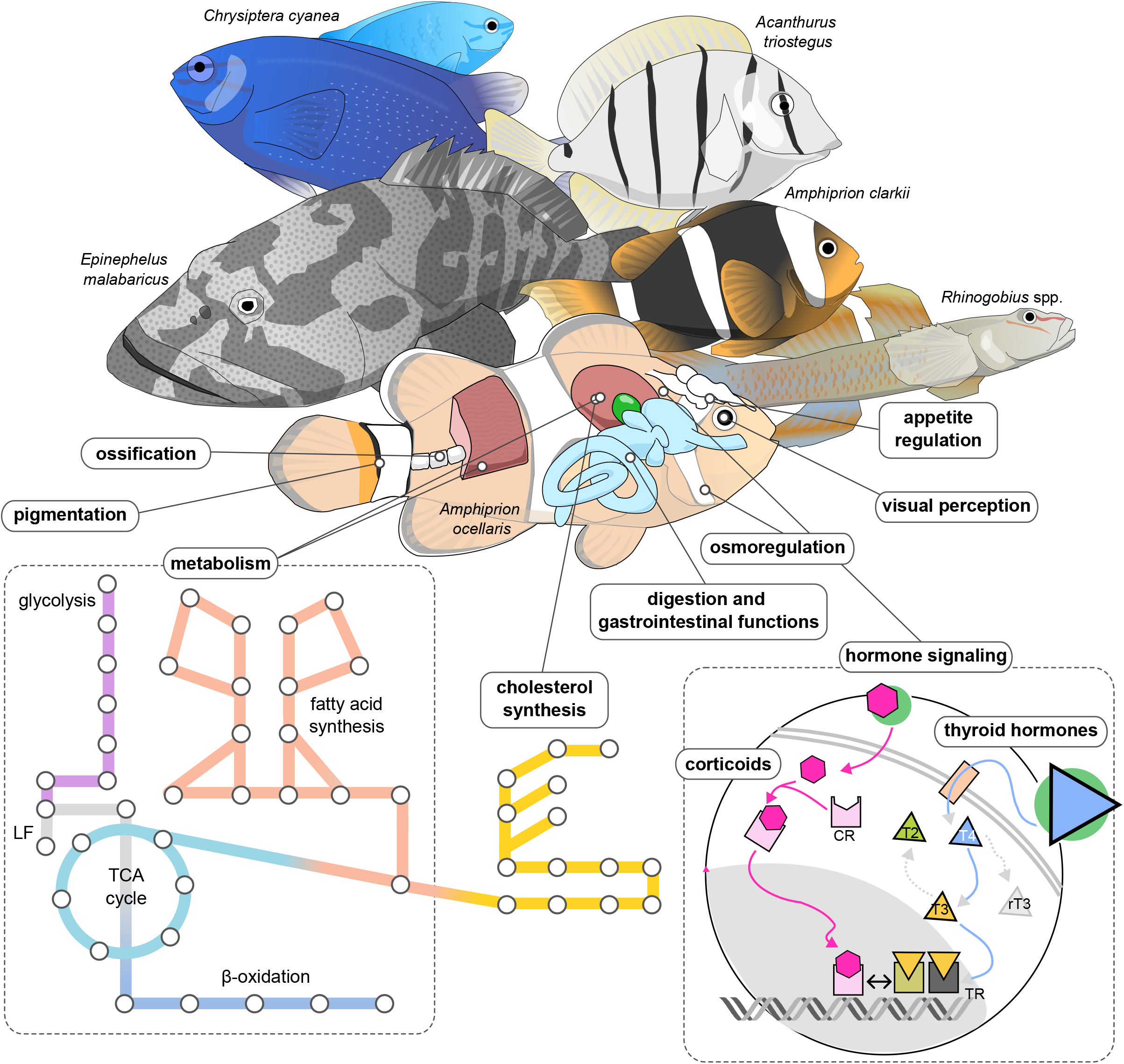
Gene repertoires for specific biological processes have been identified and validated across various species studied in our lab. These gene sets include key genes associated with hormone signaling, visual perception, appetite regulation, digestion and gastrointestinal function, osmoregulation, ossification, and pigmentation. For metabolic processes, we have selected genes that are most relevant and informative according to our data, presenting a simplified overview of key pathways, including glycolysis, lactic fermentation (LF), the TCA cycle, beta-oxidation, fatty acid synthesis, and cholesterol synthesis.

**Figure 3.**
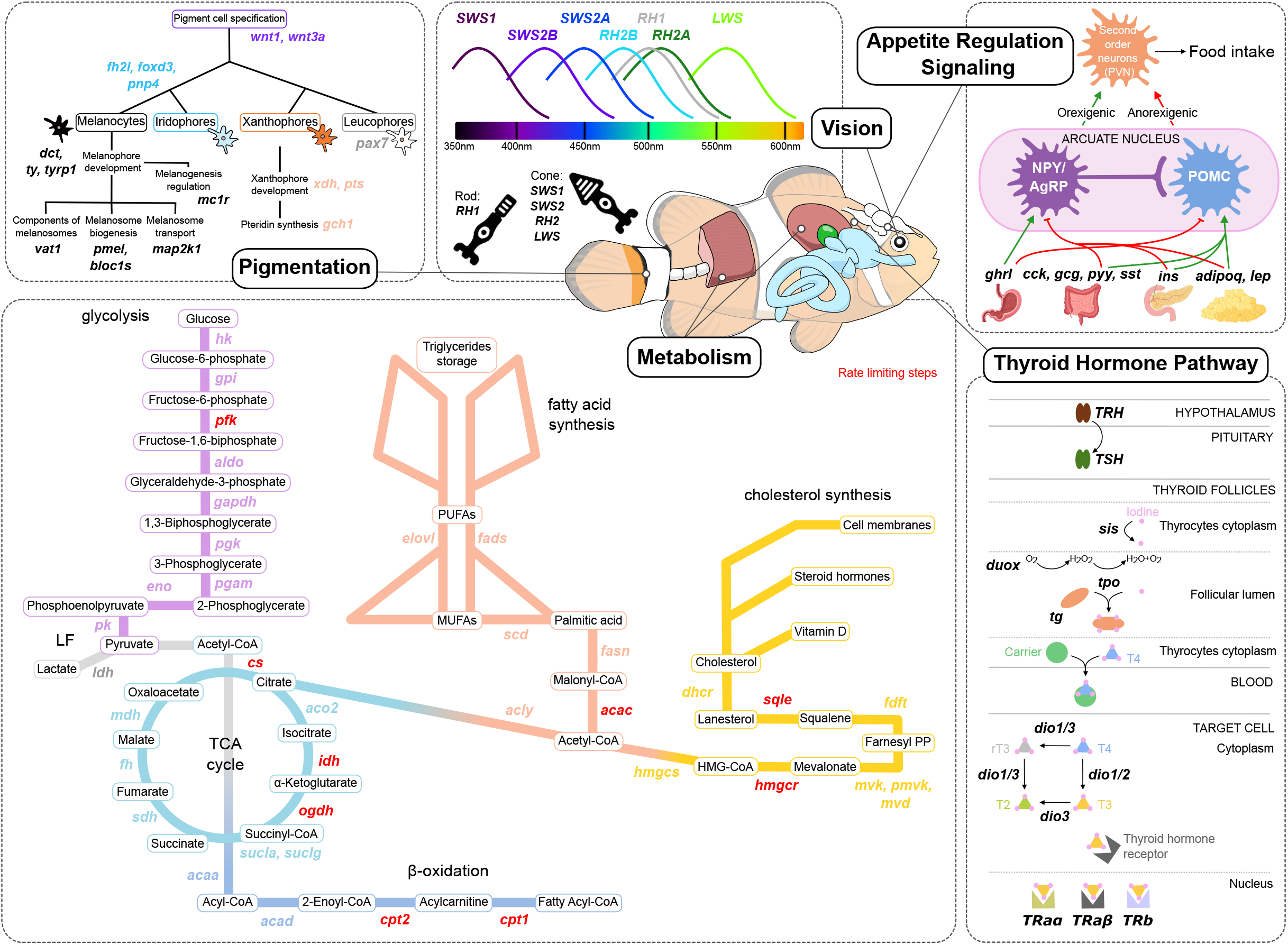
Overview of key metabolic pathways, appetite regulation, thyroid hormone signaling, pigment cells, and opsin genes featured in our gene list. Metabolism is crucial for understanding the organism’s energy needs, which are closely connected to appetite regulation and significantly influenced by thyroid hormones. Pigmentation provides insights into vital biological mechanisms and adaptations, while opsin genes reveal how fish senses and interacts with its environment. Important genes are highlighted in bold italic within their corresponding pathways.

Digestion- and gastrointestinal-related genes offer insights into feeding ecology and nutrient utilization (Solovyev et al., 2014), while visual perception genes, especially opsins and phototransduction pathways, are crucial for understanding how fish perceive and interact with their environment (Hauser & Chang, 2017; Musilova et al., 2021). Ossification genes provide a window into growth, morphology, and habitat adaptations (Campbell et al., 2021; O’Quin et al., 2015). Osmoregulation genes are essential for studying how fish maintain water and ion balance across various aquatic environments (Lema et al., 2018; Velotta et al., 2017). Lastly, we have incorporated the pigmentation genes initially identified by Lorin et al. (2018), focusing on those related to melanophores, iridophores, xanthophores, and leucophores. Pigmentation studies are crucial not only for understanding fish coloration but also as visible indicators of underlying physiological and developmental processes (Salis et al., 2021; Vissio et al., 2021). These genes often represent the “tip of the iceberg”, offering insights into less visible but equally important biological mechanisms and adaptations. Thus, by concentrating on these areas, this gene list aims to provide a comprehensive resource for transcriptomic studies investigating teleost fish responses to developmental cues, environmental changes, and ecological challenges at the molecular level (i.e., teleost fish Eco-Evo-Devo, the primary focus of our group).

## Methods

This gene list has been meticulously curated through the collaborative efforts involving specialists in various fields including fish endocrinology, metabolism, gastrointestinal function, visual ecology, skeletal development, osmoregulation, and pigmentation (Fig. 2). The criteria for selecting genes in our list were carefully considered to ensure the list’s relevance and utility. We prioritized genes based on their known or strongly predicted roles in the targeted biological processes, favoring those that show conservation across multiple teleost species as an indicator of functional importance. Preference was given to genes with experimental evidence of their function in teleost fish, rather than those solely predicted based on homology to mammalian genes. We extensively used the Zebrafish Information Network (ZFIN; http://zfin.org/) as a source of expert information to guide our gene selection. In addressing the complexity introduced by whole-genome duplication events in teleost fish, we took a nuanced approach to gene duplicates. Where functionally distinct roles have been demonstrated or strongly suggested for different paralogs, we included all relevant gene copies. To maintain consistency and facilitate cross-species comparisons, we adopted gene symbols and full gene names from the Human Gene Database (GeneCards; https://www.genecards.org/). In cases where teleost-specific gene duplicates exist, we have used Greek letter suffixes (e.g., α, β) to distinguish between paralogs. Our methodology also included a thorough phylogenetic analysis of each gene to understand their evolutionary trajectory across a diverse range of teleost species. Additionally, we retrieved species-specific identifiers for *A. ocellaris* (the original focus of our list) from the NCBI database using the publicly available genome assembly version ASM2253959v1. To ensure broad applicability, we have rigorously validated this gene list across various teleost species studied within our research group. These include several anemonefish (*Amphiprion*) species, the Sapphire devil damselfish *Chrysiptera cyanea*, the Malabar grouper *Epinephelus malabaricus*, the convict surgeonfish *Acanthurus triostegus*, and the freshwater goby *Rhinogobius* spp. (Fig. 2). Our validation process involved comprehensive analysis of RNA-seq data for each species, including tissue-specific datasets, complemented by targeted RT-qPCR experiments in selected cases.

Finally, our list is not only carefully curated, but it is also weighted to highlight key genes that serve as critical indicators or regulators of specific pathways. We have particularly emphasized pivotal genes in hormonal signaling and rate-limiting enzymes in metabolic pathways, providing researchers with important markers for these processes. Rate-limiting enzymes are significant due to their role in controlling metabolic flux through their relatively low catalysis rates. They are often extensively regulated by transcription factors and post-translational modifications, linking metabolic pathways to gene expression regulatory networks and signal transduction networks. Understanding the regulation and expression of these enzymes can provide insights into the overall metabolic state and energy demands of an organism (Fell & Cornish-Bowden, 1997). This list is the product of the collective knowledge and experience of field-specific experts across multiple disciplines of fish biology. Researchers can use this resource with confidence, knowing that each included gene has been carefully evaluated for its functional significance and relevance to teleost fish research.

### Gene List Overview

Our curated gene list comprises 739 genes that play crucial roles in key biological processes in teleost fish. These genes are organized into nine main functional categories: hormone signaling (61), metabolism (163), appetite regulation (68), digestion (121), gastrointestinal function (20), visual perception (56), ossification (24), osmoregulation (69), and pigmentation (211). The genes are listed in alphabetical order within each category and not in any specific order according to their relevance or importance. It should be noted that some genes may belong to multiple categories. We have provided detailed functional roles for most genes involved in hormone signaling and metabolic pathways, aligning with our research primary focus. However, for genes in other categories, the functional role is often left blank. This is because many genes have complex, multifaceted roles that are difficult to summarize without oversimplification.

These genes are presented in Table S1, which includes the following columns:

- Biological Process: The primary physiological or cellular function in which the gene is involved.
- Gene Symbol: The standardized short name for the gene according to GeneCards nomenclature.
- Full Gene Name: The complete scientific name of the gene according to GeneCards nomenclature.
- Gene ID (*A. ocellaris*): Identifier assigned to the false clownfish genes.
- Protein ID (*A. ocellaris*): Identifier assigned to the false clownfish proteins.
- Functional Role: A brief description of the gene’s function within the relevant biological process.

As an example of the depth and utility of our gene list, consider following the cases:

#### Case Study 1: *Citrate synthase* (*cs*)

*Citrate synthase* is a crucial enzyme in cellular metabolism, catalyzing the first step of the TCA cycle (Fisher, 2001). It catalyzes the synthesis of citrate from oxaloacetate and acetyl-CoA, a rate-limiting step that regulates the overall activity of the cycle (Fisher, 2001) (Fig. 3). It is highly conserved across species, from bacteria to humans, reflecting its fundamental role in energy production (Verschueren et al., 2019). In fish, *cs* activity is often used as a marker for mitochondrial content and oxidative capacity, making it particularly relevant in metabolic and environmental adaptation studies (Pichaud et al., 2019).

As of July 14^th^, 2024, the GO annotation for *cs* is extensive, comprising 928 terms, from which 76 terms are annotated as biological process, 81 as cellular component, and 771 as molecular function. Due to the large number of terms, they are not listed here, but the complete set can be accessed through the AmiGO 2 database (https://amigo.geneontology.org/amigo/search/ontology?q=citratesynthase). It is worth noting that 59 of these terms are currently marked as “obsolete”, indicating that they are no longer considered valid for annotating gene products. While this lengthy annotation does include terms directly referencing the TCA cycle and reflecting its regulatory role, such as GO:2000985 – positive regulation of ATP citrate synthase activity and GO:2000984 – negative regulation of ATP citrate synthase activity, it is crucial to note that identifying these specific terms among the more than 900 possible annotations can be challenging without prior knowledge. Furthermore, the vast volume of GO terms associated with *cs* makes it easy to overlook important information, particularly regarding its role as a rate-limiting enzyme and its broader implications for cellular energy metabolism. Without the expertise to select and interpret the most relevant terms, researchers might miss valuable insights.

Interestingly, when searching for KEGG pathways directly associated with *cs*, the main pathways listed is map00720 – Other carbon fixation pathways and is classified as Energy metabolism under the Metabolism module. Researchers unfamiliar with *cs* primary function could easily overlook its significance in the TCA cycle and cellular metabolism based on this classification alone. Notably, the pathway map00020 – Citrate cycle (TCA cycle), which more accurately represents the primary role of *cs*, is classified under Carbohydrate metabolism within the Metabolism module. However, this pathway might not necessarily appear in an enrichment analysis or when directly searching for “*citrate synthase*” in the KEGG Pathway database, potentially leading to an incomplete understanding of the enzyme’s function. This particular example underscores the importance of combining automated approaches with domain knowledge to ensure a comprehensive understanding of gene functions within metabolic networks.

#### Case Study 2: *Thyroglobulin* (*tg*)

*Thyroglobulin* is a critical gene in the thyroid hormone signaling pathway, encoding the precursor protein for the thyroid hormones T_4_ (thyroxine) and T_3_ (triiodothyronine) in the thyroid gland. Thyroid hormones are essential regulatory elements in the development and metabolism of all animals, with their synthesis mechanisms being evolutionary conserved (Holzer & Laudet, 2015). The overall structure of thyroglobulin is conserved from lamprey to human (Holzer et al., 2016). Its expression and function are highly conserved across vertebrates, making it an excellent model to study the molecular mechanisms of high upregulation maintenance, chromatin dynamics, and splicing kinetics (Ullrich et al., 2023).

Despite the critical role of *tg* in thyroid hormone synthesis (Fig. 3), as of July 14^th^, 2024, its annotation in the GO knowledgebase includes only three biological process GO terms: GO:1904016 – response to Thyroglobulin triiodothyronine, GO:1904017 – cellular response to Thyroglobulin triiodothyronine, and GO:0019879 – peptidyl-thyronine biosynthetic process from peptidyl-tyrosine. None of these terms directly reference the thyroid hormone pathway, which can be easily overlooked by researchers unfamiliar with hormone signaling pathways. Moreover, the ancestor charts for these GO terms do not reflect any terms related to hormone signaling. Instead, they link to broader, less specific terms such as GO:0042221 – response to chemical, GO:0070887 – cellular response to chemical stimulus, GO:0018212 – peptidyl-tyrosine modification, and GO:0044249 – cellular biosynthetic process. This case clearly demonstrates the limitations of relying solely on GO terms for functional analysis, highlighting the value of our curated gene list in providing a more comprehensive and nuanced understanding of gene function in specific biological contexts.

Similarly, its annotation in the KEGG Pathway database includes two manually drawn reference pathways: map05320 – Autoimmune thyroid disease and map04918 – Thyroid hormone synthesis. In this case, the first pathway (map05320) is likely irrelevant for fish transcriptomic studies as it references a human-specific disease and is classified as Immune disease under the Human Diseases module. On the other hand, the second pathway (map04918), which represents thyroid hormone synthesis, is more pertinent to fish research. It falls under the Endocrine System category within the Organismal Systems module and accurately represents the core process of thyroid hormone synthesis, which is largely conserved across vertebrates. However, it may have limitations for teleost studies as it does not account for fish-specific variations such as differences in regulation or the presence of duplicated genes (resulting from the teleost-specific whole genome duplication event) that might have divergent functions. While *tg* itself may not have confirmed duplicates in fish, other genes in the same pathway demonstrate this phenomenon. For example, the genes *dio3a* and *dio3b*, which encode deiodinase type III for thyroid hormone metabolism (Russo et al., 2021), are primarily found in fish, whereas mammals usually have a single *dio3* gene (Darras, 2021; Lazcano et al., 2023). This duplication, specific to teleosts allows for potentially more complex regulation of thyroid hormone activity in fish compared to mammals.

#### Case Study 3: Deiodinases (*dio1, dio2, dio3*)

The case of deiodinases presents a special challenge in genomic analyses, one that is often overlooked without specific knowledge of these enzymes. This oversight can lead to annotation issues and subsequent problems in automated analyses, potentially impacting our understanding of the thyroid hormone pathway. Deiodinases play a pivotal role in regulating thyroid hormone levels by catalyzing the activation or inactivation of T_4_ into T_3_ in various tissues (Russo et al., 2021) (Fig. 3). What makes deiodinases particularly noteworthy is their unique feature: they contain a selenocysteine residue, which is encoded by a UGA codon – typically recognized as a stop codon in most genes (Berry et al., 1991). This unusual feature can result in misannotation or incomplete annotations, leading to incorrect functional predictions or missing the full length of the protein (Kryukov et al., 2003; Lobanov et al., 2009). Failure to understand the implications of the unique selenocysteine encoding in deiodinases could lead to misinterpretations, potentially obscuring important biological insights. Thus, underscoring once again the importance of specialized knowledge to fully elucidate the complex roles and regulatory mechanisms of a specific gene.

### Practical Guide to Using this Curated Gene List

To identify orthologs across species of interest, researchers can use OrthoFinder (Emms & Kelly, 2019), a powerful tool for comparative genomics. Using the *A. ocellaris* genome publicly available in NCBI (ASM2253959v1), researchers can extract the amino acid sequences for all proteins and use them as input for OrthoFinder (Fig. 4). This approach allows for a systematic comparison of gene families and facilitates the identification of orthologs across species-specific gene duplications or losses. However, it is important to note that this is just one potential way to use the curated gene list. Researchers should feel free to adapt their approach based on their specific needs. We encourage users to explore different methodologies and tools that best suit their research objectives. This gene list can be integrated into diverse workflows and analytical pipelines, and we welcome feedback from the research community on novel ways to leverage this resource.

**Figure 4.**
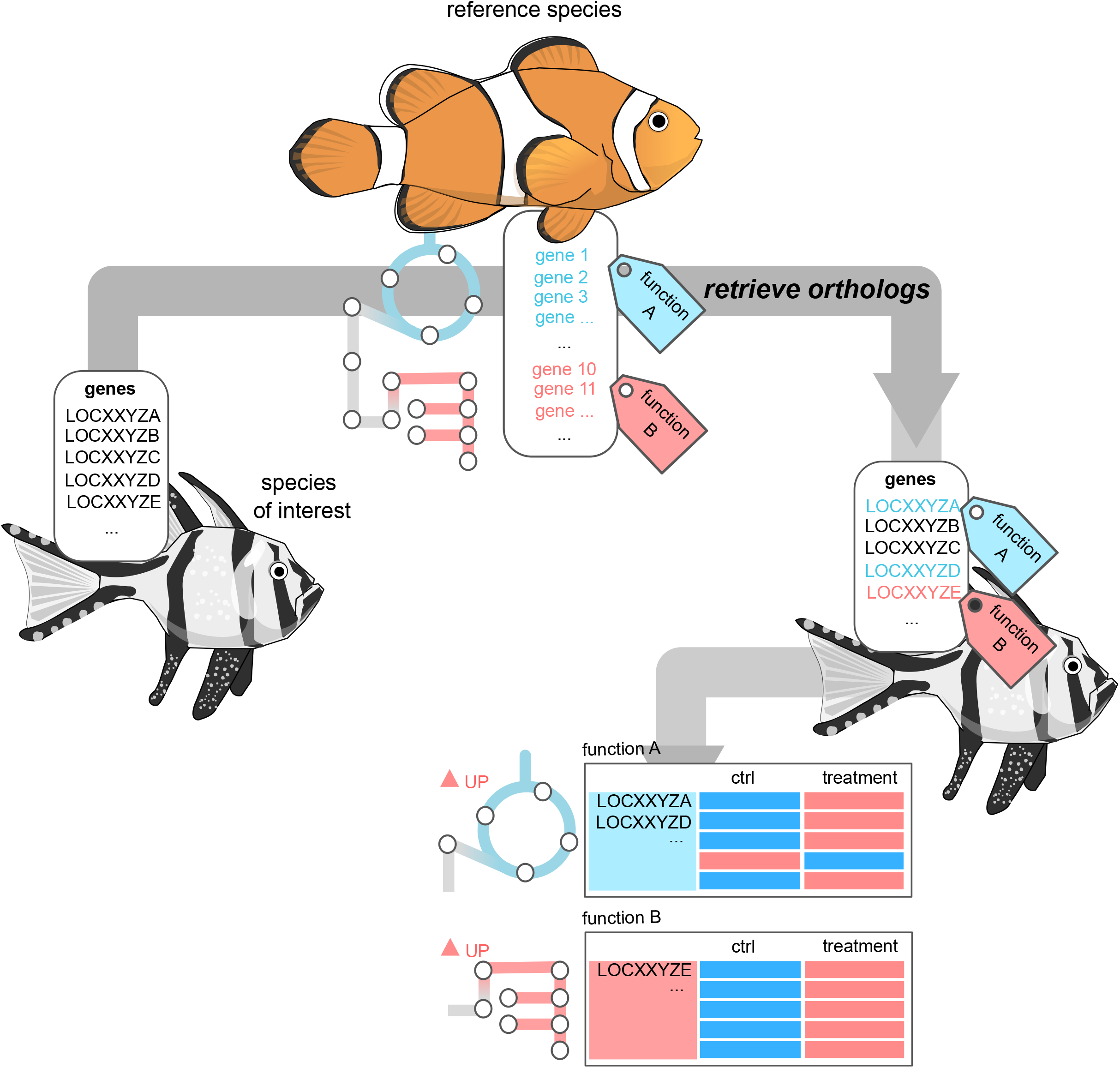
Using the false clownfish *Amphiprion ocellaris* genome (ASM2253959v1) available in NCBI, researchers can extract the amino acid sequences for all proteins and use them as input for OrthoFinder (https://github.com/davidemms/OrthoFinder) to retrieve the orthologs of the species of interest.

### Potential Applications

This gene list offers valuable applications for researchers conducting transcriptomic studies in teleost fish across several key biological processes. It can serve as a targeted panel for expression profiling, enabling quick assessment of gene activity in crucial biological processes such as hormone signaling, metabolism, digestion, nutrient absorption, visual perception, ossification, osmoregulation, and pigmentation. This comprehensive approach can be particularly useful in studies investigating developmental processes, stress responses, environmental adaptations, and physiological changes across various conditions. This list supports research on how fish respond to environmental stressors and adapt to different habitats, making it a valuable tool for ecotoxicological studies and environmental biomonitoring. Researchers can use these genes as markers to assess the impact of pollutants or climate change on fish populations, providing molecular insights into ecosystem health. For evolutionary studies, this list facilitates comparative transcriptomics across different teleost species, potentially revealing conserved and divergent gene expression patterns. It can also serve as a reference for functional enrichment analyses, helping to interpret large-scale transcriptomic data by identifying overrepresented biological processes. Researchers can use this list to design targeted RT-qPCR panels or as a reference for interpreting broader RNA-seq datasets, providing context for gene expression changes across diverse but interconnected biological systems. Overall, this curated list provides a solid foundation for hypothesis generation and targeted analysis in a wide range of teleost fish transcriptomic studies, from basic physiology to applied environmental research.

### Concluding Remarks

While this curated gene list represents a significant step forward in facilitating transcriptomic studies in teleost fish, it is important to acknowledge its limitations and context. Although the list is comprehensive in many areas, it is not exhaustive, particularly concerning metabolism-related genes. Metabolism involves a vast number of genes and complex interactions, and it would be impractical to include all of them. Instead, we have selected genes that are most relevant and informative according to our data, presenting a simplified overview of key metabolic pathways. This selection focuses on specific biological processes and pathways that we deemed most critical based on our current research and expert input. It is crucial to emphasize that this curate gene list is not intended to replace or diminish the value of established approaches such as GO term analysis or KEGG pathway mapping. These automated methods remain invaluable tools in transcriptomics, offering broad, systematic coverage of biological processes. Rather, out list of specific genes should be viewed as a complementary resource designed to enrich the biological insights that can be extracted from transcriptomic data, especially in supervised approaches where changes in specific pathways of interest are wanted to be investigated. By focusing on genes known to be particularly relevant in teleost fish biology, this list can help researchers to quickly identify and interpret key signals that might be overlooked in broader, more generalized analyses.

Future iterations of this list could benefit from incorporating additional key biological processes. For instance, expanding coverage of amino acid metabolism would provide insights into protein synthesis, energy production, and neurotransmitter synthesis (Chandel, 2021). A more comprehensive inclusion of neurotransmission-related genes would enhance the list’s utility for researchers studying fish behavior and neurobiology, allowing for more nuanced molecular phenotyping of behavioral traits (Ramírez-Calero et al., 2022). Additionally, incorporating genes involved in circadian rhythm regulation could shed light on daily activity patterns and their potential disruption by environmental stressors (Sánchez-Vázquez et al., 2019). The inclusion of aging-related genes would support research into lifespan and age-related changes in fish, an area of growing interest in vertebrate biology (Hu & Brunet, 2018). The possibilities of expansion are vast, with potential additions spanning numerous biological processes relevant to diverse research questions in teleost biology. Regular updates with newly discovered genes, refined pathway annotations, and emerging areas of research will be crucial to keep the list relevant and useful. We envision this list as a living resource, adaptable to the changing landscape of teleost fish research. The fish satellite meeting at the recent 9^th^ gathering of European Society for Evolutionary Developmental Biology (EuroEvoDevo), held in Helsinki, Finland from June 25^th^-28^th^ 2024, provided an excellent forum to discuss the importance of such collective resources. We propose to leverage the expertise of Fish Eco-Evo-Devo researchers across the globe to ensure ongoing maintenance and expansion of this list.

The significance of this gene list lies in its potential to streamline the initial stages of transcriptomic studies, as it offers a common framework to facilitate comparisons across different studies and species, potentially leading to more robust findings in teleost fish biology. By focusing on genes involved in crucial processes such as hormone signaling, metabolism, and environmental adaptation, this list supports research particularly relevant in the context of global environmental changes. The combination of our targeted approach with comprehensive GO and KEGG analyses can provide a more detailed, context-specific understanding of transcriptomic data. This synergistic use of resources allows researchers to benefit from both broad automated annotations and expert-curated gene selections, ultimately leading to richer, more biologically meaningful interpretations of transcriptomic studies in teleost fish. We believe this resource will contribute valuably to teleost fish research, providing a foundation for future work. We encourage the global research community to utilize it, offer feedback, and contribute to its ongoing development.

## Supporting information

Supplemental Table 1: Gene List

## Author Contributions

MH and VL conceptualized the idea. MH and SV compiled and finalized the gene list. Gene list revisions were conducted as follows: hormone signaling genes: NR, LB, and VL; appetite regulation and metabolic genes: NR, YG, and VL; digestion and gastrointestinal function related genes: SV; vision genes: LM; osmoregulation genes: ZC and CLN; ossification genes: EB. MH wrote the first draft of the manuscript with input from all authors. All authors read and approved the final manuscript.

## Acknowledgements

We thank the anonymous reviewers whose comments greatly improved the quality of this manuscript.

## Data Availability Statement

No quantitative data were generated. The curated gene list is presented in Table S1.

